# Using Dynamic Bayesian Optimization to Induce Desired Effects in the Presence of Motor Learning: a Simulation Study

**DOI:** 10.1101/2024.08.13.607783

**Authors:** GilHwan Kim, Haider A. Chishty, Fabrizio Sergi

## Abstract

Human-in-the-loop (HIL) optimization is a control paradigm used for tuning the control parameters of human-interacting devices while accounting for variability among individuals. A limitation of state-of-the-art HIL optimization algorithms such as Bayesian Optimization (BO) is that they assume that the relationship between control parameters and user response does not change over time. BO can be modified to account for the dynamics of the user response by implementing time into the kernel function, a method known as Dynamic Bayesian Optimization (DBO). However, it is unknown if DBO outperforms BO when the human response is characterized by models of human motor learning.

In this work, we simulated runs of HIL optimization using BO and DBO towards establishing if DBO is a suitable paradigm for HIL optimization in the presence of motor learning. Simulations were conducted assuming either purely time-dependent participant responses, or assuming that responses would arise from state-space models of motor learning capable of describing both adaptation and use-dependent learning behavior.

Statistical comparisons indicated that DBO was never inferior to BO, and, after a certain number of iterations, generally outperformed BO in convergence to optimal inputs and outputs. The number of iterations beyond which DBO was superior to BO occurred earlier when the input-output relationship of the simulated responses was more dynamic. Our results suggest that DBO may improve the performance of HIL optimization over BO when a sufficient number of iterations can be evaluated to accurately distinguish between unstructured variability (noise) and learning.

## 1. INTRODUCTION

With the advent of the era of cyber-physical systems, there has been considerable and growing interest in the use of automatic methods to measure, train, and/or augment human sensorimotor function [8, 9, 28]. In this context, multiple approaches are being explored such as: using smart activity sensors to measure repetitions, speed, accuracy, timing, and in some cases forces of human movements [5, 24]; providing participants different forms of feedback on motor performance [27]; and using active devices to physically interact with human movements [23]. The latter class of devices is of particular interest in this paper, and includes approaches such as wearable exoskeletons, functional electrical stimulation, and smart perturbation systems that are overall used to assist, augment, and improve human sensorimotor function. While the specific type of input pursued in these approaches differ widely, the underlying principle of this class of methods is to apply a physical stimulus to participants in order to induce a neuromuscular response that would modify motor coordination in a desired way, which would ultimately enable participants to accomplish the same task with less effort, or assist them to do more complex/demanding tasks with the same effort.

A specific class of devices meant to interact physically with human movement is that of wearable exoskeletons, which can assist walking by providing torques to one or multiple joints of the lower extremity [29]. These assistive torques can be defined by predefined control strategies based on position, force, or electromyography signals. How these signals are specifically used to identify the degree of torque assistance either relies on the designer/researcher intuition, or on the construction of input-output relationships based on group-level responses [22], which do not guarantee optimal results at the participant level, and can require specific, iterative tuning which can be exhaustive. Furthermore, while increased customization is possible by increasing the number of input control parameters, this modification also increases the complexity in determining the optimal set of parameters that can induce a specific response from an individual.

Human-in-the-loop (HIL) optimization was introduced to account for variability among individuals and tune control parameters in real-time [12]. Previous implementations of HIL in gait training successfully induced significant changes in participant response with different optimization methods. Recent implementations of HIL with Bayesian optimization (BO) [7, 18] have demonstrated that BO is capable of finding optimal control parameters in less time compared to other optimization methods such as covariance matrix adaptation evolution strategy (CMA-ES) [34] and gradient descent [19]. During HIL optimization, a relationship between control parameters and a corresponding cost function based on the human response is built and updated every iteration based on new measurements. Knowledge of this relationship allows the optimizer to predict future outputs for specific inputs, and to use this information to understand which inputs are more likely to optimize the cost function. However, a limitation of this technique, specifically for applications involving motor learning and rehabilitation, is that the optimizer assumes that the relationship between control parameters and cost function will not change over time or in the presence of external interventions, such as torque perturbations. This is a flawed assumption for training applications as human participants will adapt to interventions during gait training, causing their responses to the same input to change based on the history of their exposure to stimuli [25, 33].

Specifically, the human response changes to adapt to alterations in task dynamics. Previous research has introduced motor adaptation models to describe how the internal model used by the central nervous system induces changes in response during training through error-based [10, 30] and use-dependent learning (UDL) [6]. These models are generally based on a state-space formulation, where the state of the system is updated based on the history of exposure of participants to an environmental stimulus, and the state of the system modulates the input-output relationship.

These observations indicate that to accurately estimate the human response at specific points of training, the history of training, and the corresponding responses, must be considered. Therefore, to address the time varying aspects of human responses in training with HIL optimization, an optimizer must be capable of considering time as a component in the human response model.

A modified BO method (dynamic Bayesian optimization - DBO) has previously been developed and validated to address time-varying systems, i.e., systems with explicit time-dependency in the input-output equation [1, 2, 21, 26, 35]. Past studies have shown that DBO performed better compared to existing optimization methods such as BO, CMA-ES, and particle swarm optimization in finding the optimal input of predefined dynamic problems such as 6-hump Camelback, Griewank, and Shekel functions [26]. While DBO has not yet been implemented nor simulated for HIL optimization, it may improve upon BO as the human response can be treated as a general time-dependent system in complete absence of a model of human neuromotor adaptation. However, the applicability of DBO has not been tested in scenarios constructed to emulate human adaptation, and so its feasibility for application in HIL optimization remains untested.

In this work, we implemented DBO in HIL simulations capturing key features of the human response to robot intervention during training. The virtual human participant response was constructed to reflect dynamic responses due to time-dependency or arising from state-space models of motor learning. State-space models were based on a model of the human response that incorporates features of adaptation and use-dependent learning. For benchmarking of DBO, convergence speed and accuracy of the virtual HIL optimization are compared between DBO and BO.

## 2. METHODS

### 2.1 Dynamic Bayesian Optimization

BO is a probabilistic approach to finding the optimal input value that induces a desired outcome from a system based on data that is observed with measurement error [3]. Given a set of observations and a predefined covariance function accounting for the statistical relationship between changes in input values and changes in the outputs, BO will construct a Gaussian process (GP) model which estimates the responses for unobserved values of the input. Based on the GP model, BO uses an acquisition function to determine the input to be tested in the subsequent iteration: this acquisition function provides higher scores for inputs that result in good predictions (exploitation) or high prediction uncertainty (exploration). To inhibit overexploitation, which could lead to the optimizer being trapped in local minima, an additional parameter known as the exploration-exploitation ratio (e-ratio) is implemented. For a given GP model, the the model variance at input 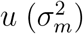 is defined as the sum of the variance of cost function 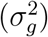 and additional noise (*σ*^2^) as:

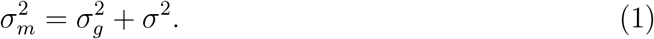

The BO algorithm will flag overexploitation if the next test input *u*_*next*_ satisfies

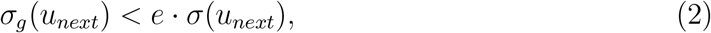

where *e* is the e-ratio. Thus, when higher e-ratios are used, the optimizer is more likely to flag inputs as overexploitation, and therefore select new inputs that offer greater uncertainty (exploration). If overexploitation is still detected, hyperparameters in the covariance function are adjusted, and the GP model is reconstructed.

In classical BO, observation time is not considered as it is assumed that the system is stable over time. Therefore, repetitions of the same input values are expected to generate similar observations. To account for the fact that a system response may change with time, as is the case of neuromotor adaptation, BO needs to be modified.

To account for the time-varying response of individuals to a set of inputs, the covariance function *k*(*u, u*^*′*^) between two sets of input parameters *u* and *u*^*′*^ can be augmented by introducing the time variable *t* as *k*((*u, t*), (*u*^*′*^, *t*^*′*^)), where *t* and *t*^*′*^ are the time instances when inputs *u* and *u*^*′*^ are applied, respectively. Previous research simplified this covariance function by assuming separability to describe the time-component of covariance function [2, 13]. Therefore, the overall covariance function can be described as multiplication of a static component of the covariance function *k*_*u*_ and a dynamic component *k*_*t*_, as

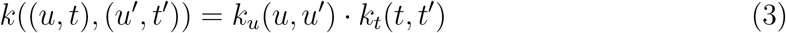

#### 2.1.1 Static Component of Covariance Function

The static component of the covariance function *k*_*u*_ used in a Gaussian process model is only determined by input parameters *u*. In this work, we defined the static (i.e., input-dependent) component of the covariance function based on the most commonly used squared exponential covariance function:

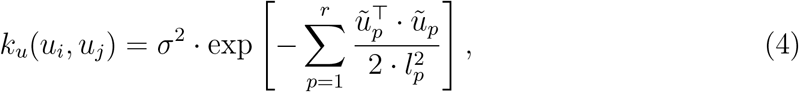

where *u*_*i*_ and *u*_*j*_ are *r*-dimensional inputs at arbitrary time points *i* and *j* respectively, *r* is the number of input/control parameters, and *ũ*_*p*_ is the difference between the *p*-th input parameter of two input parameter arrays *u*_*i*_ and *u*_*j*_ (*u*_*ip*_ *− u*_*jp*_). *l*_*p*_ is the *p*-th length scale hyperparameter, which influences the function’s sensitivity to differences in parameter inputs, and *σ*^2^ is the measurement variance.

#### 2.1.2 Dynamic Component of Covariance Function

Considering the idea that two measurements collected with a larger time difference will be less related to each other than two measurements collected with a smaller time difference, the dynamic component of the covariance function *k*_*t*_ is assumed to have a lower value as the time difference between two data points increases. In this work, we defined the dynamic (i.e., time-dependent) component of the covariance function *k*_*t*_ between two time points *t* and *t*^*′*^ consistently with previous research [2] as:

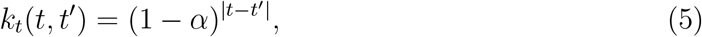

where *α ∈* [0, 1) a hyper-parameter. Therefore, the overall covariance function used in this work is

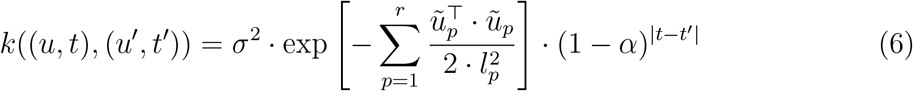

### 2.2 Optimizer Testing on a Model with Explicit Time Dependence

To assess the effect of incorporating the dynamic component in the covariance function used in Gaussian process modeling for optimization, both versions of the optimizer were compared using two virtual HIL simulations with predefined time-varying responses: 1) a system with explicit time dependence and 2) a set of state-space models of neuromotor adaptation.

Both processes used Expected Improvement as the acquisition method based on previous research demonstrating its effectiveness in convergence with a similar problem involving walking biomechanics [4, 17, 31].

#### 2.2.1 Explicit Time Dependence Model

To include explicit dependency on time or iteration number in the input-output relationship, we defined a process as a zero-order system, where the output would change with iteration number in three ways: 1) the optimal output would change with time as dictated by a decaying oscillating exponential function; 2) the output would deviate from its optimal value quadratically based on the distance between the current input and the optimal input; and 3) the optimal input would itself change with time. The input (*u*)-output (*y*) equation for this system at each iteration (*i*) is defined as:

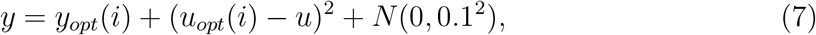

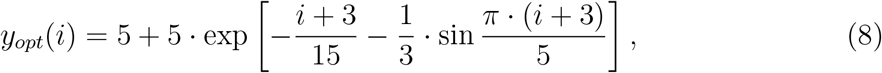

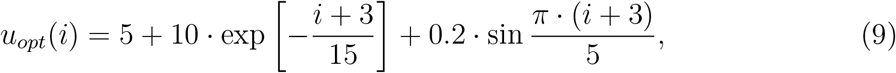

where *i* is the iteration number, and *N* (0, 0.1^2^) is observation noise sampled from a normal distribution with zero-mean and standard deviation equal to 0.1. *u*_*opt*_(*i*) is the optimal input at the *i*-th iteration, and *y*_*opt*_(*i*) is the corresponding optimal response of the system at the *i*-th iteration. The optimal value of input and response are shown in Fig. 1. This type of system was selected due to its explicit time-dependence: the associated response to a newer optimal input would be lower (i.e., more optimal) than a past optimal input. Therefore, an ideal optimizer should rely more heavily on a recent estimation of the optimal input to achieve the desired response (*y* = *y*_*opt*_).

**Figure 1.**
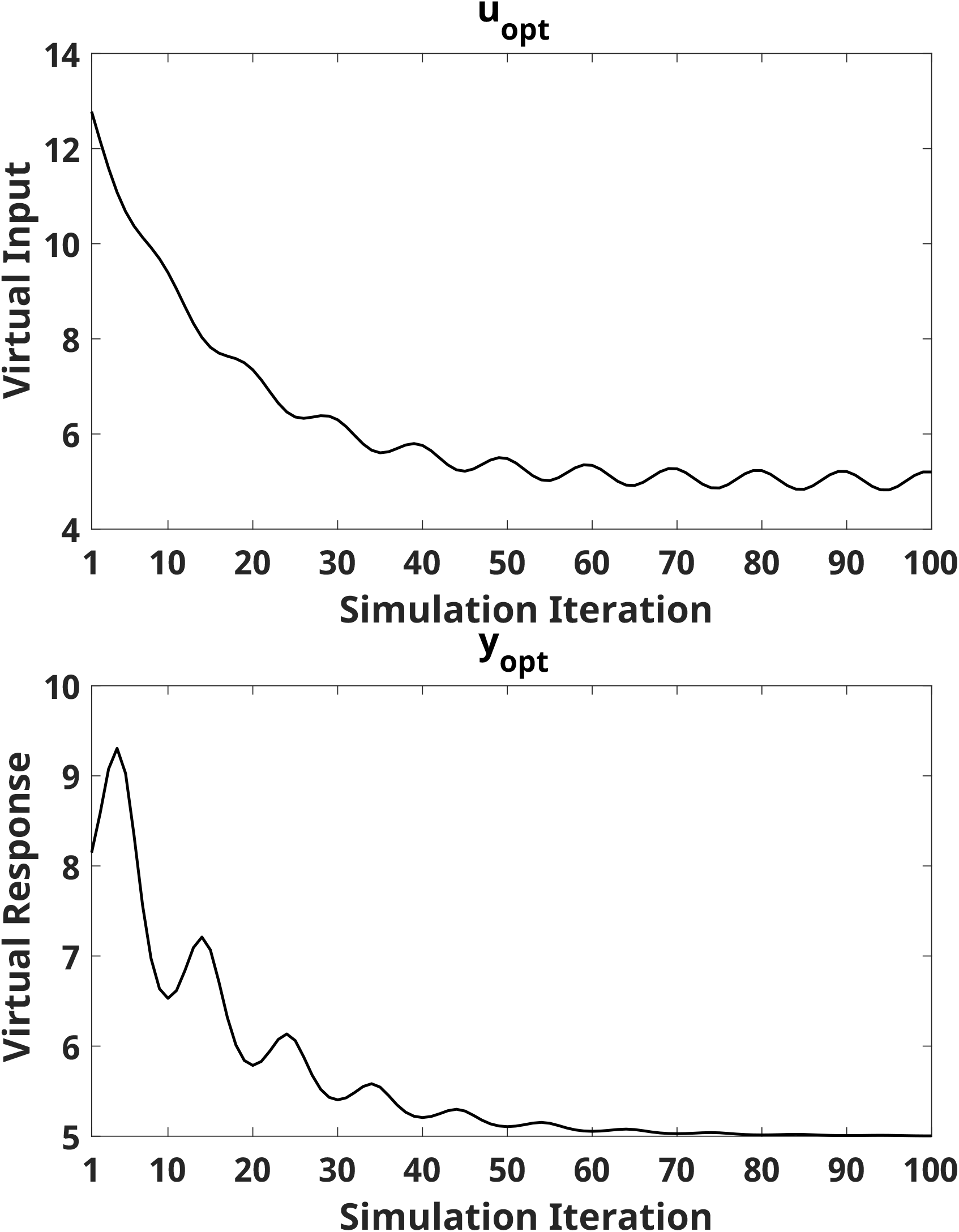
(Top) Optimal input value *u*_*opt*_ that results in the optimal response *y*_*opt*_ at every iteration (*x*-axis). (Bottom) Optimal response *y*_*opt*_ at every iteration. *x*-axis indicates simulation iteration number, all occurring after three initial inputs.

#### 2.2.2 Simulation Method

Both DBO and BO were implemented to simulate a virtual HIL experiment when the input at each iteration was applied to the model, and the output was calculated using Eq. (7)-(9). Both methods were implemented with different e-ratios within the range [0.2:0.2:1]. A higher e-ratio resulted in an optimizer that favored exploration rather than exploitation.

Based on the Gaussian process model generated at each iteration, each optimizer made an estimation *u*_*estim*_(*i*) of the optimal input *u*_*opt*_(*i*) that minimized the response *y*(*i*) from the system at each iteration. Obviously, the optimizer did not have any knowledge of the structure of the system; however, knowledge of the true system equation was used for benchmarking of the optimizers. Specifically, we quantified the *output error* as the absolute deviation between the measured response *y*(*i*) and the optimal response *y*_*opt*_(*i*), and the *input error* as the absolute deviation between the applied input *u*(*i*), and the optimal input at that iteration *u*_*opt*_(*i*). Moreover, the accuracy of the model estimated by the optimizers about the time-varying relationship between optimal input and optimal output was assessed by quantifying the *input estimation error* as the deviation between the true optimal input *u*_*opt*_(*i*), and the estimated optimal input *u*_*estim*_(*i*), as well as the *output estimation error*, as the absolute deviation between the true optimal output *y*_*opt*_(*i*) and the output estimated as the response associated with the estimated optimal input *f* (*u*_*estim*_(*i*)).

Simulations were repeated 100 times following three initial inputs, selected randomly. For each repetition, the same set of three initial input values and corresponding responses were used in both DBO and BO to start HIL optimization. Each simulation was run for 100 iterations, not including the initial three inputs. To ensure wide enough input range during optimization, bilateral saturation limits |*u*| *<* 15 were included.

### 2.3 Optimizer Testing on a State-space Model of Learning

Motor adaptation models describe how the internal model used by our central nervous system to control our movements changes in response to changes in task dynamics. To describe how the central nervous system refines these models, error-based learning models are widely used [15]. In an error-based learning model, the internal model is updated by two processes: a feed-forward signal predicted from internal dynamics, and an error-based component that updates the model based on the perceived error. Errors are often calculated based on the difference between the actual and planned movement or force. To address use-dependent effects in the human response, where individuals often rely on past movements to decide the current movement, a UDL model was previously introduced [6].

While a plethora of motor learning models have been proposed [14, 20], most of the proposed motor learning models share the fact that they are iterative learning models, where the state of the system is updated based on the history of exposure of participants to an environmental stimulus. For these systems, the relationship between the stimulus *u* and the output *y* is not explicitly time-dependent, but is rather dependent on the value of a certain number of states, that may or may not be directly observable given a characterization of the learner dynamics. A general form of such a state-space system is provided below:

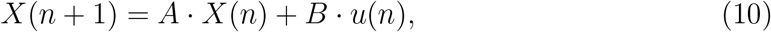

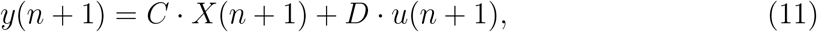

where *X*(*n*) = [*x*_1_(*n*), …, *x*_*p*_(*n*)]^*⊤*^, *A ∈* ℝ^*p×p*^, *B ∈*ℝ^*p×r*^, *C ∈*ℝ^*q×p*^, and *D ∈* R^*q×r*^, *u ∈*ℝ^*r*^, and *y ∈*ℝ^*q*^ .

#### 2.3.1 Modified Use-dependent Learning Model

Even for a linear system, the number of open parameters scales dramatically with the order *p* of the state space equations, making exhaustive analysis of all model configurations intractable. However, our own previous research indicates that a modified UDL model, an instance of a 2^nd^ order linear state model of motor adaptation, was a sufficiently accurate and parsimonious model for describing the effects of exoskeleton inputs on propulsion mechanics [16]. This model offers a more accurate representation of human response than the time-varying response tested in Sec. 2.2 as generated responses are based not only on inputs but also on past responses.

The modified UDL model is formulated by introducing an update equation for the reference state *x*_0_ in the existing UDL model, to account for the possibility of after-effects of training. In this model, the state *x*_0_ is a reference movement in absence of any stimulus, the state *x* is the planned movement, the input *u* is the applied stimulus (i.e. torque/force), and the output *y* is the participant response. All state parameters (*x*_0_, *x, u*, and *y*) are updated at every iteration *n* as:

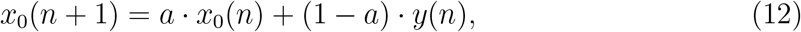

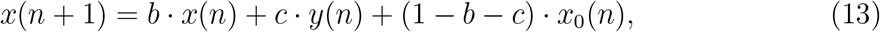

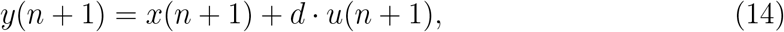

where *a* is the retention parameter for the reference state bounded on [0 1], *b* is the previous movement retention parameter, *c* is the use-dependent learning term, and *d* determines the sensitivity of the human response changes to robot-applied input *u*(*n*). Based on the previous movement *y*(*n −* 1), the reference movement *x*_0_(*n*) is updated, affecting the planned movement *x*(*n*) at the current iteration *n*. The standard state-space form for the modified UDL model with state vector *X*(*n*) = [*x*_0_(*n*), *x*(*n*)]^*T*^ is

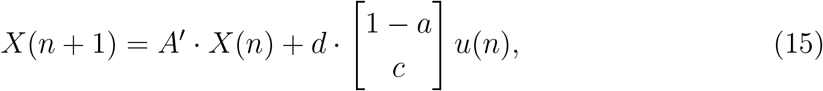

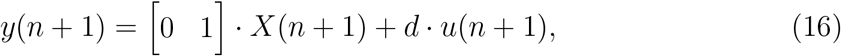

where 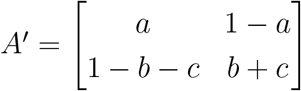. Four parameters (*a, b, c*, and *d*) define this modified UDL model.

Different combinations of parameters *a, b, c*, and *d* allow the modified UDL model to account for different amounts of adaptation, use-dependent learning, and remaining after-effects following training. For example, we define a positive learning model as a model that continuously increases the response in the presence of a positive stimulus. In contrast, a negative learning model continuously decreases response in the presence of a positive stimulus. Two different learning dynamics were selected for the positive and negative learning models to evaluate the advantages of DBO at different learning rates. Specifically, we expected performance difference between DBO and BO to be higher when the system response exhibited higher dynamics i.e., greater changes in response to continuous stimuli. Fast learning models were designed to generate changes in response greater than 100% between the initial and final responses when the same continuous inputs were exerted over 200 iterations. In contrast, slow learning models were designed to make less than 50% changes between the initial and final responses under the same condition. These four models, as well as two others, were selected for virtual human-in-the-loop simulation experiments, and can be seen in Fig. 2, where responses in the presence of 200 iterations of continuous stimulus and 100 iterations without stimulus are shown (as well as an initial 100 iterations of no stimulus).

**Figure 2.**
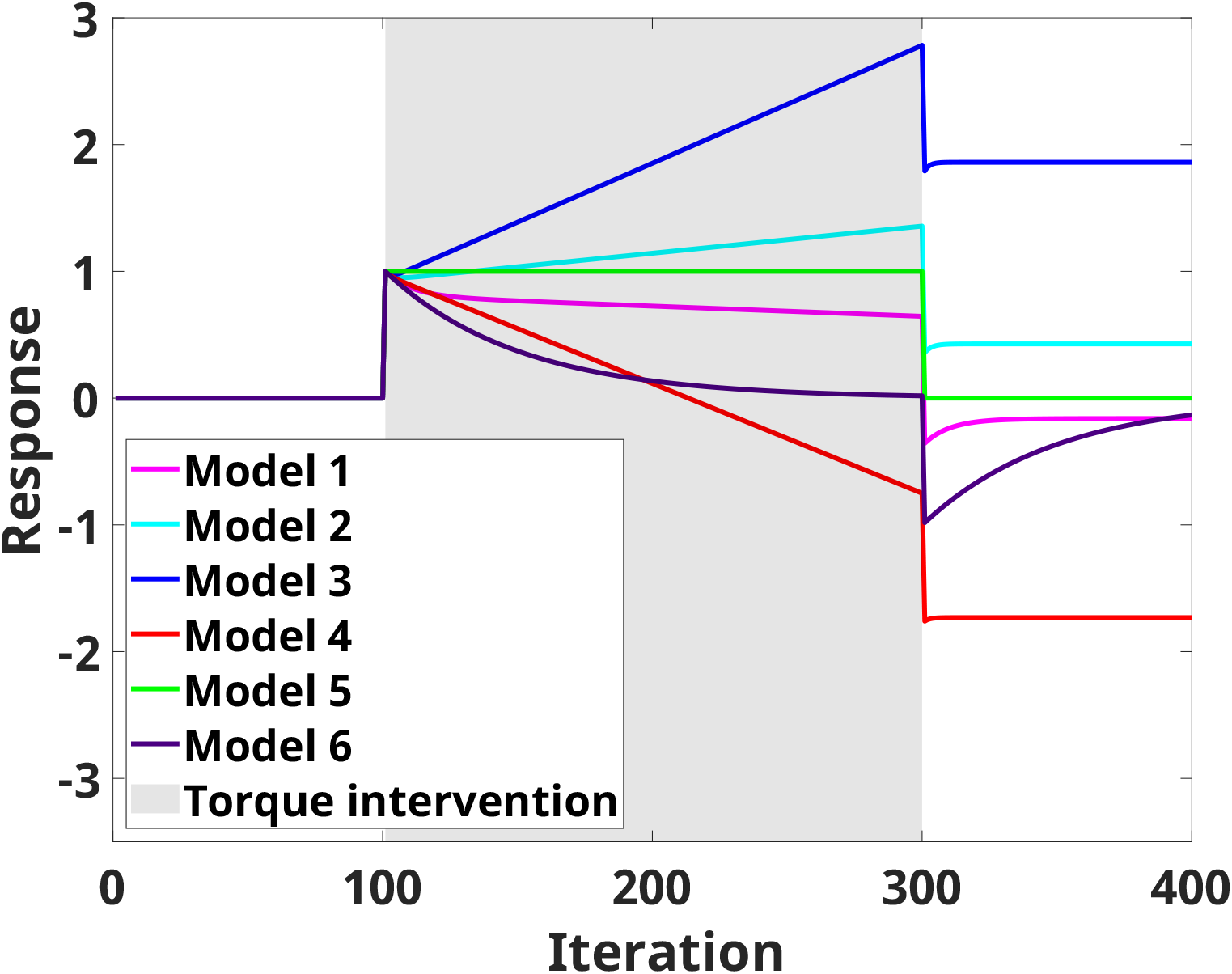
Response (*y*) from different modified UDL models when 100 iterations of null input, followed by 200 iterations of constant input (*u*=1), and 100 iterations of zero input are applied (Model 1: slow negative learning, Model 2: slow positive learning, Model 3: fast positive learning, Model 4: fast negative learning, Model 5: no learning and no adaptation, Model 6: only adaptation).

A specific instance of the modified UDL model highlighting its behavior is shown in Fig. 3. The states *x* and *x*_0_ as well as the output *y* increase when positive inputs *u* are continuously applied (positive learning); in contrast, response *y* decreases when negative inputs *u* are continuously applied. When no inputs are applied, state and response values are maintained.

**Figure 3.**
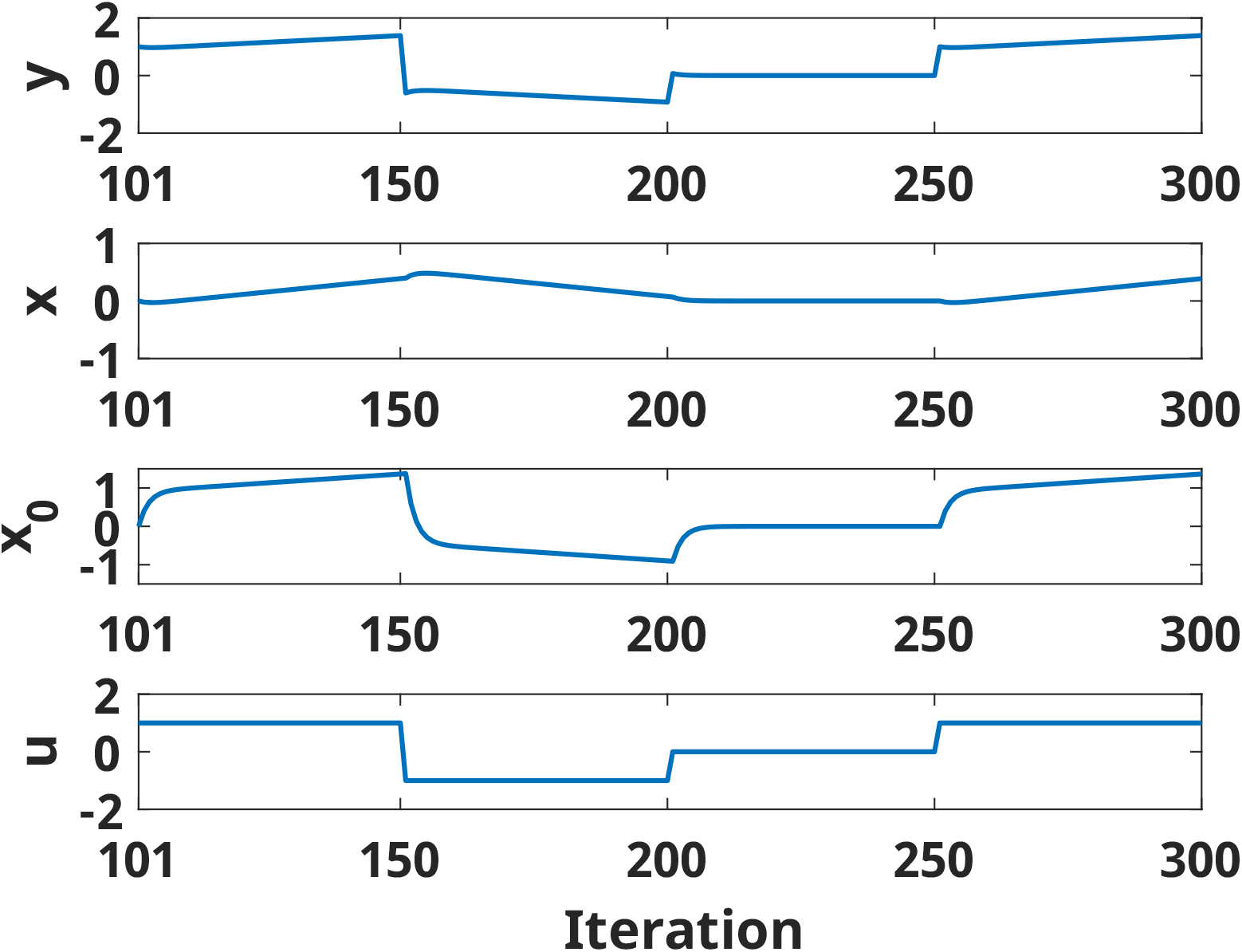
States (*x* and *x*_0_) and output (*y*) of the modified UDL model with positive learning at different iterations when different inputs *u* are applied (50 iterations of 1, 50 iterations of −1, 50 iterations of 0, and 50 iterations of 1). The states and output change even when a constant (or null) input is applied

#### 2.3.2 Simulation Method

HIL simulations were conducted using the state-space models described in Sec. 2.3.1 as virtual human responses. Responses were influenced by the history of the inner states (*x* and *x*_0_), response (*y*), and input (*u*). However, during simulation, only the response was provided to the optimizer; i.e., the optimizers were unaware of the state-space structure of the system. Therefore, to induce desired responses during simulation, the optimizers needed to estimate the relationship between input and system responses by properly weighing the relevance of previous observations, subject to the effects of hidden states *x*(*n*) and *x*_0_(*n*).

Simulations were conducted using both versions of the optimizer, using e-ratios that provided the best convergence and estimations of optimal inputs/outputs for the HIL simulations described in Sec. 2.2 (i.e., 0.2 and 0.4). Each simulation was performed over 50 repetitions; all repetitions had the same three initial inputs (stimulus) - selected randomly - consistent across optimizers. 200 iterations were simulated via optimization to achieve a response of *y* = 1 (i.e., to minimize the cost function *g* = (*y −* 1)^2^). Observation noise was added to the output *y* as a random variable sampled from *N* (0, 0.1^2^), i.e., a normal distribution with zero-mean and 0.1 standard deviation.

HIL simulations with both optimizers were performed assuming that the virtual human response would result from the six model types described in Sec. 2.3.1 and Fig. 2. Parameter values for each model were decided upon using participant-specific fitting results from previous research [22], and the capability of describing a wide range of adaptation behaviors. Model parameters and the objective responses are seen in Table 1. Bilateral saturation limits |*u*| *<* 3 were included when simulating Models 1, 2, and 5; for Models 3, 4, and 6 saturation limits were instead |*u*| *<* 10.

**Table 1.** Coefficient Parameter Values and Objective Response of Neuromotor Adaptation Models.

| | $a$ | $b$ | $c$ | $d$ | $y_{obj}$ |
| --- | --- | --- | --- | --- | --- |
| Model 1 | 0.9202 | 1.001 | -0.02 | 1 | 1 |
| Model 2 | 0.7 | 0.9977 | -0.02 | 1 | 1 |
| Model 3 | 0.6 | 0.99 | -0.02 | 1 | 1 |
| Model 4 | 0.6 | 1.089 | -0.02 | 1 | 1 |
| Model 5 | 0 | 1 | 0 | 1 | 1 |
| Model 6 | 1 | 1 | -0.02 | 1 | 1 |

At each iteration *i*, the response from the modified UDL model depends on the system states *x*(*i−*1) and *x*_0_(*i−*1), on the output *y*(*i−*1), and input *u*(*i*). Therefore, the optimal input *u*_*opt*_(*i*) that achieves the target response *y*_*obj*_ at iteration *i* is calculated as:

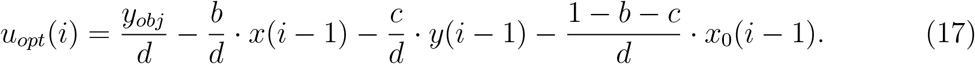

Similar to the simulation using the model with explicit time dependency, *input error, output error, input estimation error*, and *output estimation error* were collected from simulation results to validate whether each optimizer was able to dynamically adjust their input based on the observed learning effects to achieve the desired constant response during simulation.

### 2.4 Statistical Analysis

For both optimization problems, the simulation outcomes *input error, output error, input estimation error, output estimation error*, as defined previously (see Table 2) were subject to statistical analysis to establish whether they were different across optimizers. Specifically, once normality was confirmed via Kolmogorov-Smirnov tests, we constructed iteration-specific two-way ANOVA models with factors Optimizer type (two levels: BO and DBO), and Optimizer E-Ratio (two settings: 0.2 and 0.4), and ran the analysis of a full factorial two-way ANOVA model separately at each iteration. We reconducted all ANOVAs using the “fitlm” MATLAB function, to account for unequal variances, to confirm if this assumption impacted the results of our two-way ANOVAs. In presence of significant effects or interactions, post-hoc t tests were implemented. Given the nature of this dataset fully resulting from numerical simulations and with arbitrary sample size, model effects were reported at an uncorrected significance level *p*_*unc*_ *<* 0.05, for each iteration.

**Table 2.**
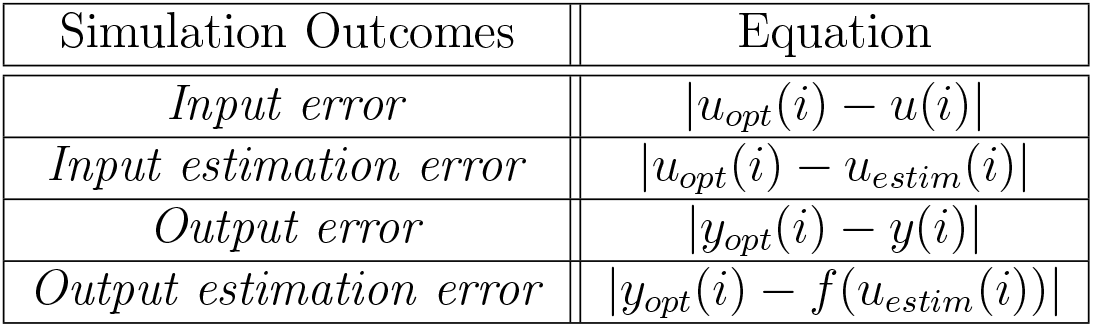
Simulation outcomes and description.

To facilitate analysis and comparison of effects across outcomes, we classify the outcomes into two types: one set, describing the accuracy by which the optimizer estimates the optimal inputs/outputs (estimation outcomes, i.e., *input estimation error* and *output estimation error*), and one set describing the accuracy of the current inputs and outputs relative to the currently optimal ones (implementation outcomes, i.e., *input error* and *output error*).

## 3. RESULTS

### 3.1 Optimizer Testing on a Model with Explicit Time Dependence

Results from simulations using an explicitly time-dependent system as the predefined response are seen in Fig. 4, where absolute differences between the optimal input and output (*u*_*opt*_ and *y*_*opt*_ respectively) are compared to those implemented (*u* or *y*) or those estimated (*u*_*estim*_ or *y*_*estim*_) by each optimizer. All data presented here were confirmed normally distributed by Kolmogorov-Smirnov tests, and ANOVA results were confirmed to have not been impacted the by equal variance assumption. As shown by the shaded regions in Fig. 4, the two-way-ANOVA demonstrated that DBO is continuously superior to BO after a certain number of iterations: these significant differences occur following iterations 12, 11, 13, and 13 for *input error, input estimation error, output error*, and *output estimation error* respectively.

**Figure 4.**
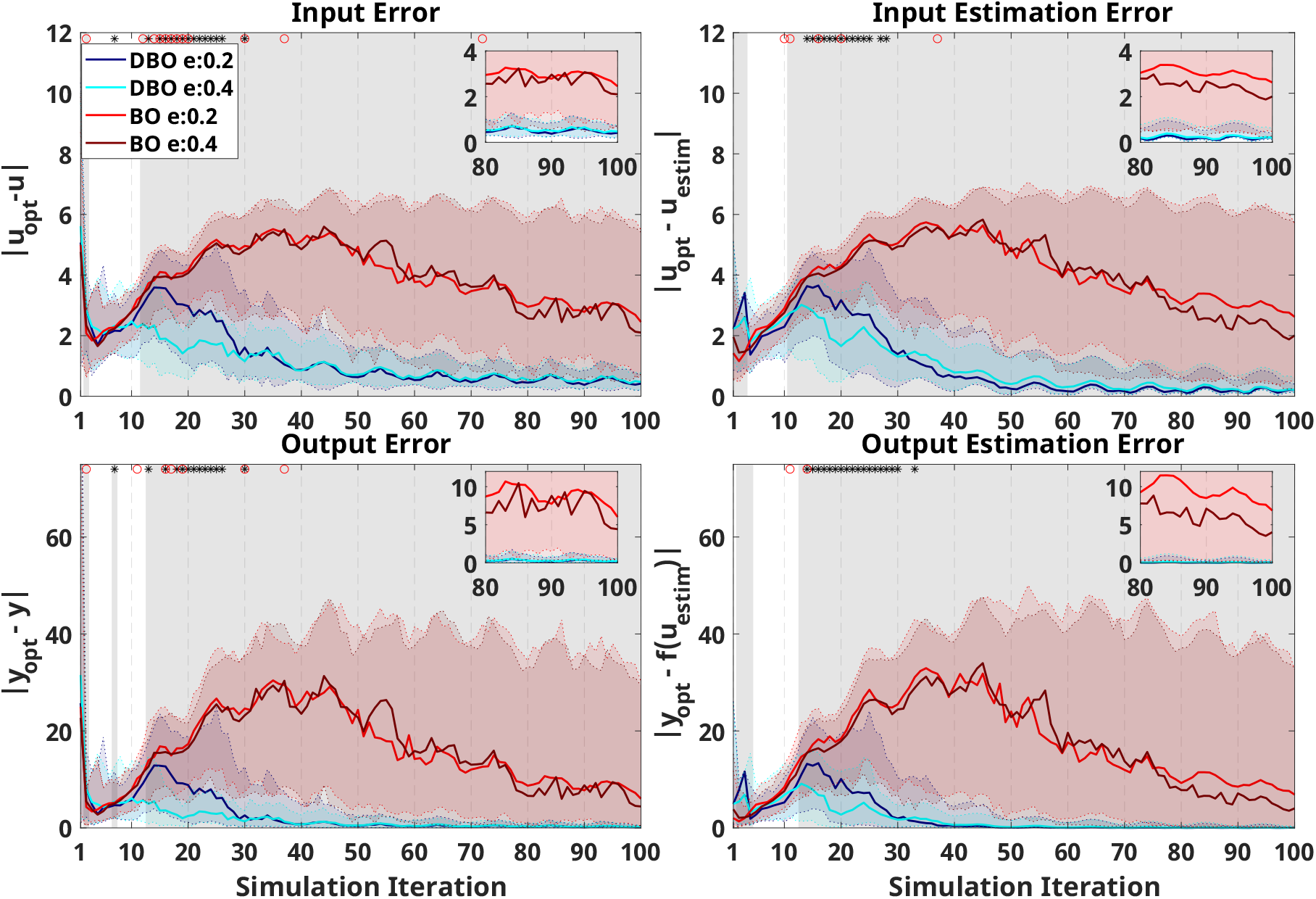
Optimizer performance on the selected model with explicit time dependence. The *x*-axis indicates the number of iterations during optimization (i.e., after three random inputs are initially applied). In all plots, the blue/cyan and red/crimson indicate simulation results using DBO with e-ratio of 0.2/0.4, and BO with e-ratio of 0.2/0.4, respectively. Solid lines indicate median and shaded regions of same color extend from the 20^th^ to the 80^th^ percentile across 100 repetitions at each iteration. Optimizer performance is quantified based on *input error* (top left), *input estimation error* (top right), *output error* (bottom left), and *output estimation error* (bottom right). A zoomed-in insert is provided in each panel to help visualize differences in the last 20 iterations. The gray shaded region indicates the presence of significant differences (*p*_*unc*_ *<* 0.05) between simulation results using different optimizers at each iteration. Asterisks at the top of each figure indicate a significant effect of the e-ratio, and red circles indicate a significant interaction between e-ratio and the optimizer on the outcome at each iteration.

Significant effects of e-ratio, or of its interaction with optimizer type, were present in the early stages of training, with e-ratios of 0.4 outperforming those of 0.2, primarily for the DBO optimizer. Post-hoc analyses over this significance region (estimated to be between iterations 10-30) indeed confirmed that there was no significant effect of e-ratio within the BO optimizer (only one iteration showed significantly different *input error* and *output error*), while for DBO, e-ratio had a significant effect in 81% (*input error*), 76% (*input estimation error*), 71% (*output error*), and 86% (*output estimation error*) of the iterations tested between iterations 10 to 30. In these cases, DBO results with an e-ratio of 0.4 were superior to e-ratio of 0.2 in all cases except two iterations on *input error*, one iteration on *input estimation error*, and two iterations on *output estimation error*. Except one iteration for the *input error* outcome, no significant effects of e-ratio were measured past iteration 37.

Example results from a single trial are seen in Fig. 5. As can be seen, DBO outperformed BO over all outcomes, with both implemented and estimated inputs of DBO with an e-ratio of 0.2 converging to the optimal values soon after iteration 30, and with DBO with an e-ratio of 0.4 converging with some error. BO was unable to capture changes in the system, although BO with an e-ratio of 0.4 did trend towards to the optimal near the end of simulation.

**Figure 5.**
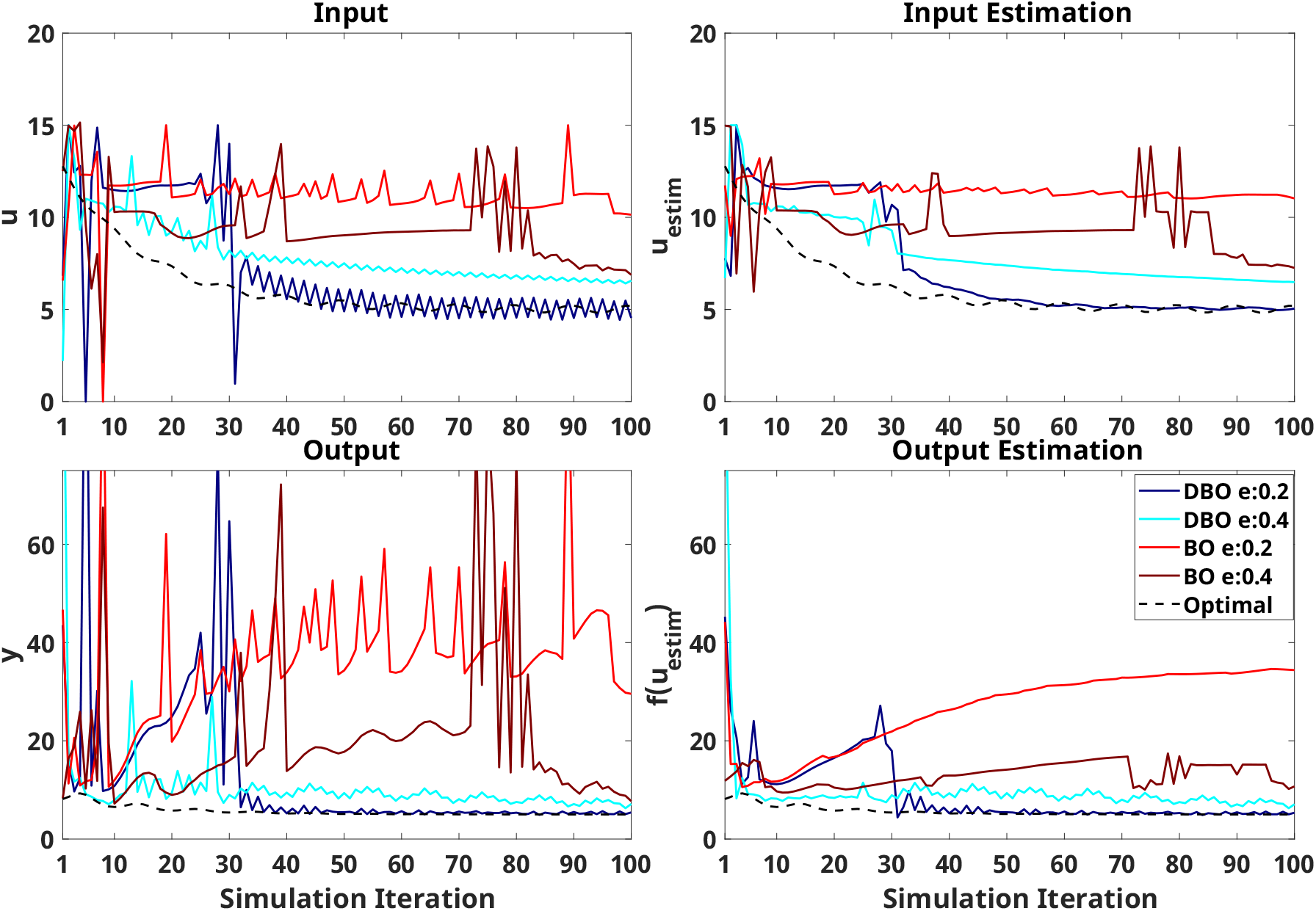
Simulation results of single trial using selected model with explicit time dependence as the virtual human response. In all plots, the blue/cyan and red/crimson indicate simulation results using DBO with e-ratio of 0.2/0.4, and BO with e-ratio of 0.2/0.4, respectively. Black dash lines indicate optimal input or output at each iteration.

### 3.2 Optimizer Testing on a State-space Model of Learning

Results of the virtual human-in-the-loop simulations interacting with the state-space model of learning are reported in Fig. 6, 7, S2, S3, S4 and S5 for the six tested model types. As above, all data presented were confirmed normally distributed by Kolmogorov-Smirnov tests, and ANOVA results were confirmed to have not been impacted the by equal variance assumption. Results associated with Models 3 and 4 (fast positive and negative learning) are discussed below in greater detail as these models have a faster dynamics, and thus DBO is expected to more accurately model the input-output relationship.

**Figure 6.**
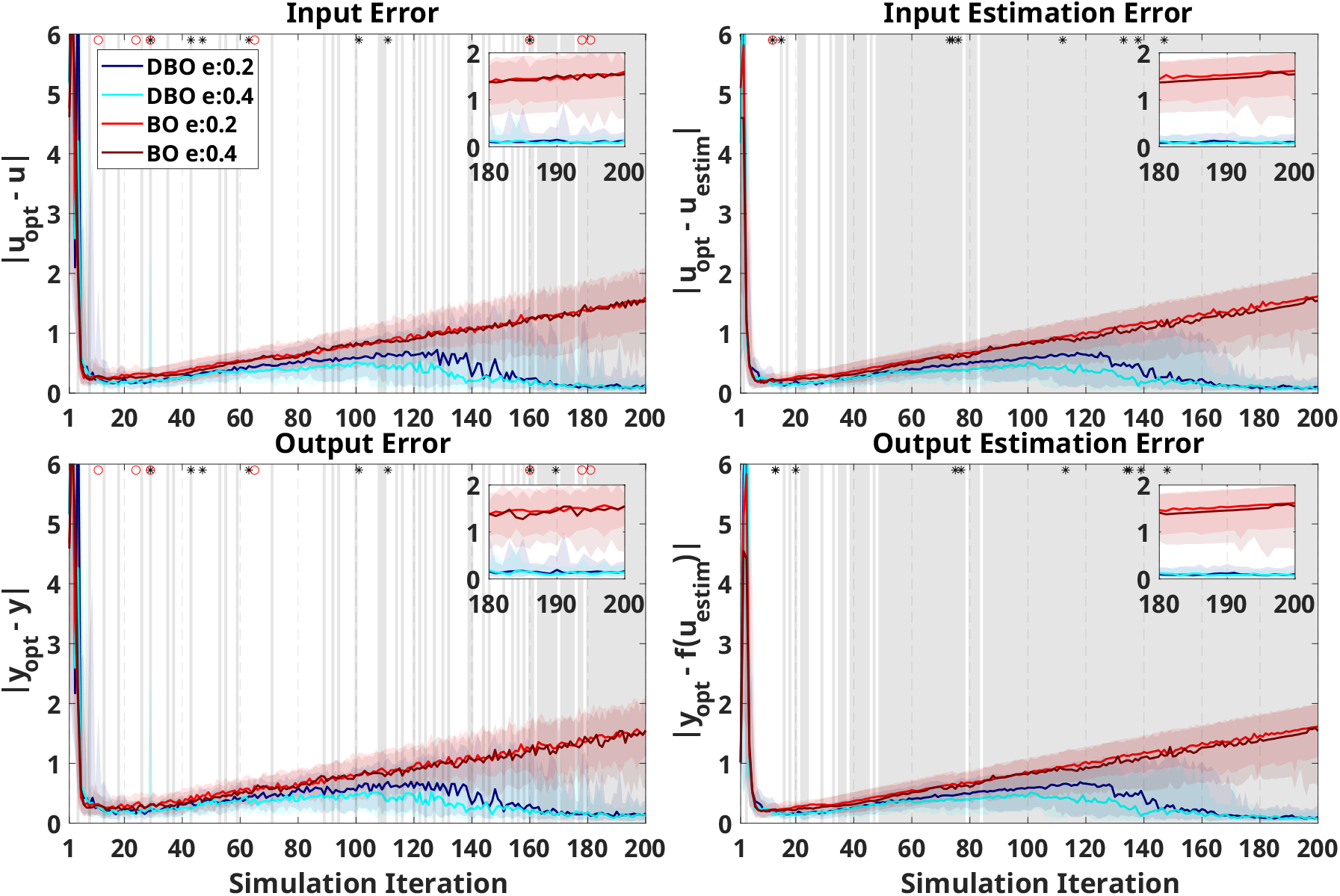
Virtual HIL simulation results using UDL model with positive learning (Model 3) as the virtual human response. The *x*-axis indicates the number of iterations during optimization (i.e., after three random inputs are initially applied). In all plots, the blue/cyan and red/crimson indicate simulation results using DBO with e-ratio of 0.2/0.4, and BO with e-ratio of 0.2/0.4, respectively. Solid lines indicate median and shaded regions of same color extend from the 20^th^ to the 80^th^ percentile across 50 repetitions at each iteration. Optimizer performance is quantified based on *input error* (top left), *input estimation error* (top right), *output error* (bottom left), and *output estimation error* (bottom right). A zoomed-in insert is provided in each panel to help visualize differences in the last 20 iterations. The gray shaded region indicates the presence of significant differences (*p*_*unc*_ *<* 0.05) between simulation results using different optimizers at each iteration. Asterisks at the top of each figure indicate a significant effect of the e-ratio, and red circles indicate a significant interaction between e-ratio and the optimizer on the outcome at each iteration.

**Figure 7.**
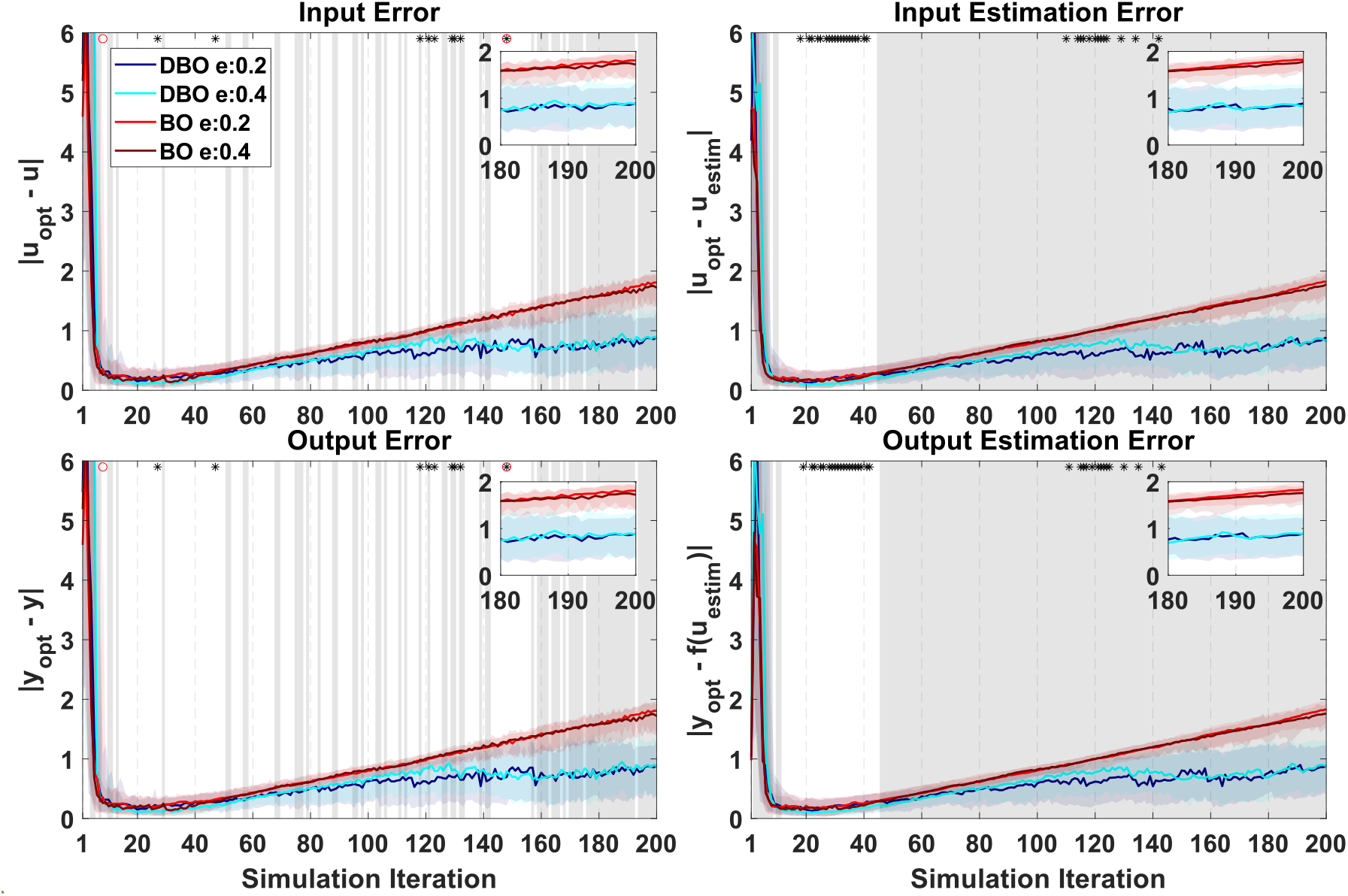
Virtual HIL simulation results using UDL model with negative learning (model 4) as virtual human response.

Overall, results from the two fast learning models indicate that statistical differences associated with different optimizers were not as continuous beyond a certain number of iterations as in the explicitly time-dependent model, but a clear pattern emerged where DBO optimizers were superior to BO as iteration number progressed. Similarly, differences due to the e-ratio were prominent in the early stages of training, but some occurrences of e-ratio effects were also observed in later stages of training.

Specifically, for Model 3 (Fig. 6), a two-way ANOVA demonstrated that DBO is superior to BO in 90% of the iterations past iteration 12 for *input estimation error* and *output estimation error*. For Model 4 (Fig. 7), the two-way ANOVA revealed that DBO is superior to BO in 100% of iterations past iteration number 44.

Less continuous trends are observed for the implemented outcomes, where DBO only achieves superiority in 80% of iterations following iterations 147 and 149 for *input error* and *output error*, respectively in Model 3 (Fig. 6), and following iteration number 152 for both outcomes in Model 4 (Fig. 7). Moreover, the patterns of errors are distinct between the two optimizers: while errors seem to grow indefinitely for BO in both models, average DBO errors decrease after a certain number of iterations for Model 3, while they appear to remain roughly constant for Model 4.

Significance due to e-ratio or its interaction with optimizer type was very sparse within Model 3 (Fig. 6). There were only a few iterations showing a significant effect of e-ratio within the BO optimizer, and all below the false-positive rate of 0.05 (10 iterations per outcome). Similarly for the DBO optimizer, the largest number of iterations where a significant effects of e-ratio was measured was for *input error*, where the optimizer with e-ratio of 0.4 significantly outperformed the one with 0.2 in 7 out of 10 iterations.

In Model 4, significance due to e-ratio was sparse for outcomes *input error* and *output error*, but almost continuous in certain iteration regions for *input estimation error* and *output estimation error*. Post-hoc analyses showed that there were only two iterations for both *input error* and *output error* that showed significant effect of e-ratio within the BO optimizer. Within the DBO optimizer, 7 iterations for both *input error* and *output error* and 28 iterations for both *input estimation error* and *output estimation error* showed a significant effect of e-ratio, the latter above the false positive rate of this test. *Input estimation error* showed significant e-ratio effects in 59% of iterations between iterations 10 and 50, with 0.4 outperforming 0.2, and 10% of iterations between iterations 110 and 150, with 0.2 outperforming 0.4 instead. For *output estimation error*, 56% of iterations between iteration 10 and 50 exhibited significant difference due to e-ratio, where 0.4 outperformed 0.2, and 12% of iterations between 110 and 150 exhibited this significance, where 0.2 outperformed 0.4.

In slow learning models (Models 1 and 2), effects of optimizer type were qualitatively similar as those measured in the fast learning models, but the improvements achieved with DBO were quantitatively smaller. Specifically, no simulation outcomes showed consistent significant difference prior to iteration 150; however, DBO was superior to BO continuously for outcomes (*input estimation error* and *output estimation error*) at the end of simulation (beyond iteration 150).

Simulation outcomes using only adaptation (Model 6) showed significantly greater DBO performance for estimation outcomes in most iterations from iteration 54 to the end.

Simulations assuming a constant response (Model 5) did not show significant difference for optimizer type or e-ratio for all outcomes in most iterations (number of iterations where a significant effect of optimizer type was detected was below the false-positive rate of 0.05). In all analyses conducted for Models 1, 2, 5, and 6, the effect of e-ratio was minor, and significant at a rate below the false-positive rate of the statistical test.

Fig. 8 shows single trial results using a negative learning model (Model 4). Results from this trial indicate more exploration from DBO in the later phase of simulation (Fig. 8 - Top and Bottom left); further, results show that DBO is able to estimate optimal inputs/outputs better (Fig. 8 - Top and Bottom right).

**Figure 8.**
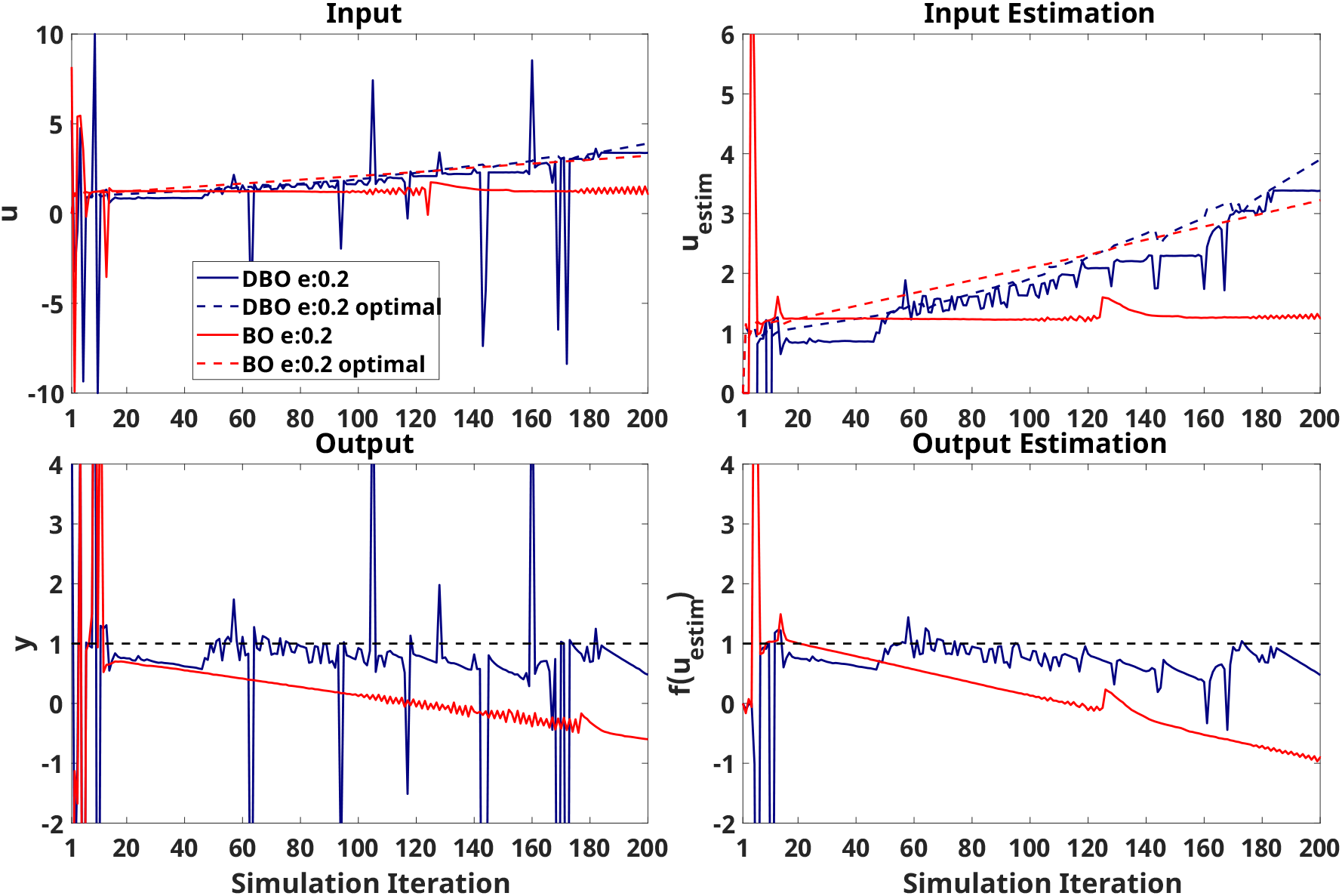
Simulation results of single trial using UDL model with negative learning (Model 4) as virtual human response. Blue and red solid line indicate result from DBO and BO respectively. Dashed line is optimal input or output during simulation.

## 4. DISCUSSION

Towards the goal of establishing if dynamic Bayesian optimization is a suitable paradigm for human-in-the-loop optimization in the presence of a learning agent (such as human participants adapting to interventions during gait training), DBO was used in HIL simulations where virtual participant responses were constructed to reflect dynamic responses due to time dependency in the model, or due to lack of information about state variables provided to the optimizer. Results of these simulations were compared to simulations done using conventional BO, and convergence speed and accuracy were compared between the two optimizers.

The state-space model representing neuromotor adaptation and learning, the modified use-dependent and error-based learning (UDL) model, used in the HIL optimization simulation to generate virtual human responses was developed and validated in the context of explaining the evolution of propulsion mechanics in response to exoskeleton assistance [16]. This model best captured the measured human responses compared to alternative neuromotor models. By adjusting a few parameters, the modified UDL model can describe different forms of learning and adaptation, including the four types of responses (positive/negative, learning/adaptation) identified in previous research [22]. Therefore, this model enables characterization of individual responses to robotic intervention during training and prediction of the after-effects of training.

The state-space models designed to represent different levels of adaptation and learning (Fig. 2) required the input to continuously increase or decrease to maintain the same output. Thus, the optimal input needed to elicit the desired response will continue to change as the iterations progress. As the internal state values and corresponding optimal inputs continually change, the system exhibits a non-stationary response. Since the optimizer has access only to past input–output data, an optimizer that performs better than another can be interpreted as having a more accurate estimation of the input–output relationship, leading to more precise tracking of the desired outcome. However, due to the lack of direct information about the internal state values, the optimizer must inherently handle a non-stationary system.

Across the time-varying responses described in this paper used for HIL simulations, DBO consistently performed better than BO if the response was highly dynamic (such as in a model with explicit time-dependence, or in the case of state-space models of fast neuromotor adaptation). This is characterized by significantly lower absolute differences when comparing both the implemented and estimated outcomes at the final stages of simulation with the those from BO; further, two-ways ANOVAs demonstrated that DBO converged to mostly continuous significant improvement over BO after a certain number of iterations, occurring fairly early when the virtual response was based on an explicitly time-dependent model (iterations 11 to 13). In the case of the fast state-space models, this significant difference again occurred early for the estimated outcomes *input estimation error* and *output estimation error* (occurring at iterations 12 and 44 for Models 3 and 4 respectively).

The differences in optimizer performances can be attributed to the evaluations associated with new observations: specifically, when a new input-output observation is acquired, BO will adjust uncertainty to take into account the new observation along with all past observations; in contrast, DBO will aim to separately fit two sources of variance between measurements, one due to measurement/process noise, and one due to the fact that responses are measured at different time points and are naturally bound to be different due to that.

This rationale is consistent with the results of simulations using other state-space models. Using the faster, 1^*st*^ order, purely adaptation state-space system (Model 6), DBO superiority is achieved after a small number of iterations. Meanwhile, in the slower dynamic models (Models 1 and 2), DBO superiority is achieved much later. Finally, in the essentially stationary constant response model (Model 5), no DBO superiority is ever achieved, but the dynamic optimizer performs just as well as its BO counterpart.

Regardless of the difference in nature of an explicit time-dependent response and fast state-space models with hidden states, there are a number of similarities in simulation results. Namely, all simulations experience heavy transience early on, as the optimal input changes drastically during early states of simulation. Following this early transience, DBO begins to improve over BO, specifically in estimation outcomes where significant differences appear early and remain mostly continuous over the rest of simulation. For both of the fast state-space models, this continuous significance for implemented outcomes begins later, as the optimizers continue to explore unseen parameters in attempts to maintain or improve estimation accuracy.

Results of HIL simulations using BO do differ based on which system is used for the virtual human response, as outcome errors do eventually decrease with iteration when the response is based on the explicit time-dependent system - this is due to the implemented decay of the optimal output with iteration, which drives BO to value newer observations, and the relatively stationary optimal input-output relationship in the latter half of simulation, which allows BO to begin to converge to the optimal input. In contrast, BO is unable to capture the non-stationarity of the state-space responses, and instead fixates on optimal inputs found early on in simulation, causing outcome errors to increase with iteration.

DBO and BO were tested with different e-ratios in all model cases: since the e-ratio heavily affects how the optimizers select subsequent inputs, we hypothesized that optimizers with different e-ratios will result in different simulation results. No significant differences in simulation results were found using different e-ratios in BO; in contrast, different e-ratios affected the accuracy of simulated DBO runs, primarily in the early iterations (i.e., before iteration 30). The statistical comparisons described in detail were only conducted using e-ratios of 0.2 and 0.4, as they demonstrated DBO’s best performance when simulating using the explicitly time-dependent model as shown in Fig. S1.

As described above, HIL simulations using faster, time-varying responses indicated that DBO generally outperformed BO as simulation iterations proceed, across outcomes; however, as indicated previously, statistical significance of the between-optimizer comparisons were not always consistent across outcomes. Specifically, implementation and estimation outcomes are affected differently by the ratios of exploration vs. exploitation that the optimizers maintain at different stages of optimization. From single-trial visualizations (Fig. 8), it is possible to observe that DBO exhibits greater levels of exploration compared to BO, especially later through the simulation. As such, while estimation outcomes typically indicated continuous DBO superiority at some point during simulation, this continuity was not as strong in implementation outcomes; in fact, DBO’s greater exploration resulted in greater variability of these outcomes, thus reducing the signal-to-noise ratio of between-optimizer comparisons.

The exact number of iterations needed to observe a substantial difference between DBO and BO will depend on the speed of the learning process, as evidenced by fast and slow dynamic models exhibiting this difference at different points of simulation. Other aspects, such as the signal-to-noise ratio of the observations, possible process errors, the range of inputs, and the number of input parameters may also affect how many iterations are needed. These application-specific variables need to be considered accurately when selecting whether to use DBO or BO for a specific problem. To this aim, it is important to note that our analysis did not identify any condition where DBO was inferior to BO, with the exception of only the very first few iterations in some of the runs.

Practical considerations need be made to translate these simulation-based outcomes into practice. Fatigue is obviously a critical concern for the real implementation of HIL optimization algorithms, as fatigue would set an upper bound on the iteration number, requiring optimizers to converge to desired solution in a quite limited amount of time. This could be addressed by dividing training over multiple sessions or providing participants with sufficient breaks to avoid the negative effects of fatigue. Additionally, the computational time required for each iteration may also affect the total experiment duration, as it increases the number of strides taken per iteration. The computation time primarily increases with the amount of past data and the number of control inputs considered in the optimization. This issue can be addressed by using a dedicated, high-performance computer exclusively for the optimization process. Under more constrained conditions, such as those involving excessive measurement noise or stringent time limitations, alternative or complementary algorithms to DBO and BO may need to be explored to reduce computational demands. Previous work showed that HIL optimization can improve outcomes such as metabolic cost compared to baseline by 17.4% in only 20 iterations [7] - our analysis does not indicate that DBO is superior to BO for in this early iteration range. Instead, we propose that DBO may be advantageous in HIL optimization cases where greater numbers of iterations are achievable, such as those that target biomechanical outcomes that are quantifiable in less time than metabolic outcomes, and those where learning effects may be more relevant.

Furthermore, there are possible consequences of DBO that may not be fully captured in our simulations: as visible in single trial results (Fig. 8), DBO leverages exploration more than BO in latter portions of the simulation - such drastic exploration results greater variance of sequential inputs applied to the user. From the motor learning literature, we know that variance in environmental dynamics - be it task variability or external stimulus - affects human adaptation behavior [25]. For example, participants exposed to a highly unpredictable perturbation will use co-contraction to increase joint stiffness and reject a perturbation, while participants exposed to a predictable stimulus may be able to respond by adapting their forward model to take advantage of the changed task dynamics [11, 32]. Such input-variance dependent behavior is not captured by our state-space model, and thus it is possible that such highly variable sequential inputs may affect user performance in ways that we are unable to model via our simulations.

In conclusion, we evaluated the capability of DBO in implementing HIL optimization towards inducing desired changes in participant responses. Both DBO and BO estimated system responses and attempted to converge to optimal inputs. Seven different models including one time-explicit system and six neuromotor adaptation models were tested in virtual HIL simulations. Overall, results indicate that DBO performs better than BO in estimating optimal input and subsequent system response, and may improve the performance of HIL optimization over BO when a sufficient number of iterations can be evaluated to accurately distinguish between unstructured variability and learning.

## 7. Supplementary Materials

**Figure S1:**
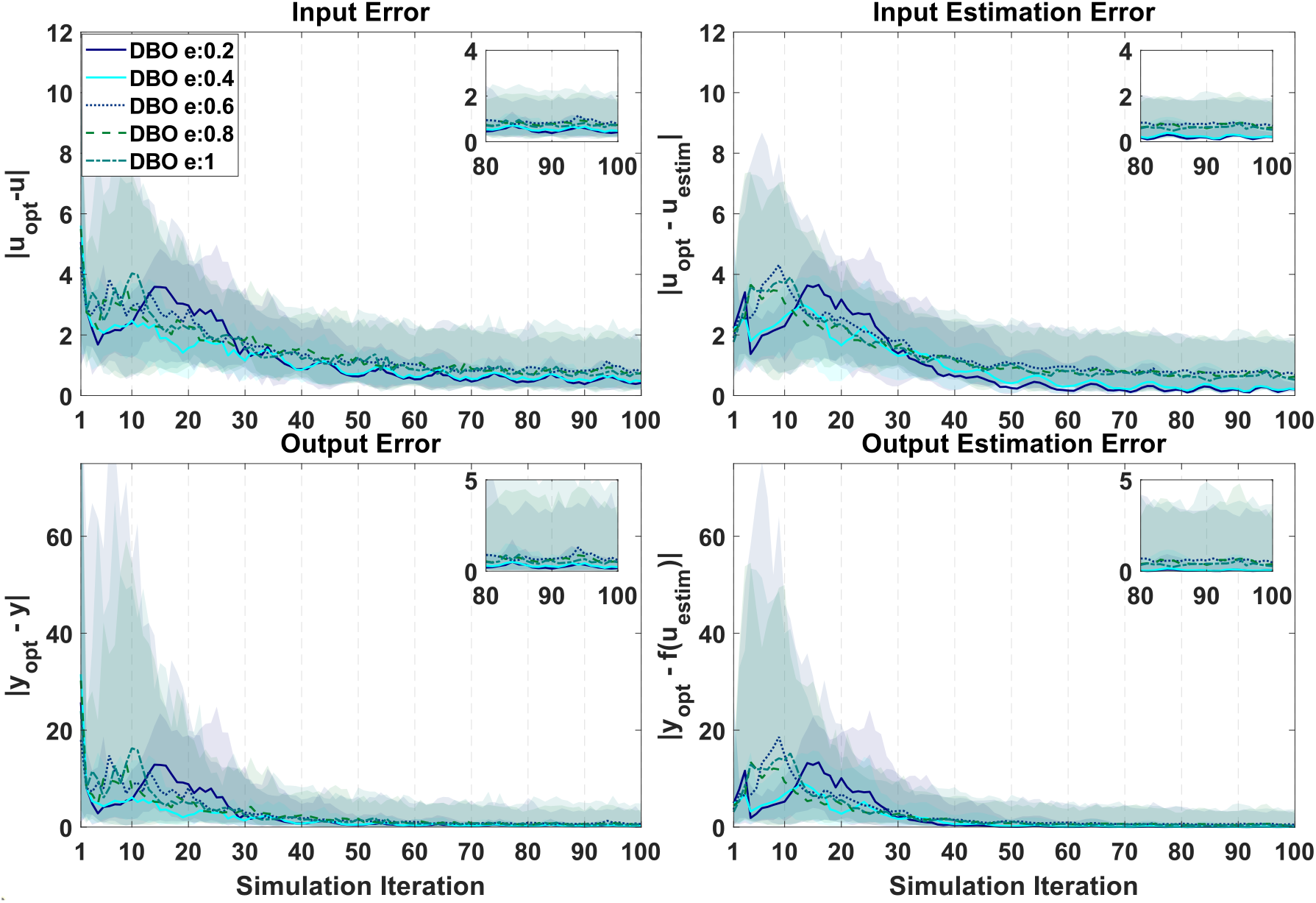
DBO optimizer performance on the selected model with explicit time dependence. The *x*-axis indicates the number of iterations during optimization (i.e., after three random inputs are initially applied). In all plots, solid lines indicate median and shaded regions of same color extend from the 20^th^ to the 80^th^ percentile across 100 repetitions at each iteration. Optimizer performance is quantified based on *input error* (top left), *input estimation error* (top right), *output error* (bottom left), and *output estimation error* (bottom right). An zoomed-in insert is provided in each panel to help visualize differences in the last 20 iterations.

**Figure S2:**
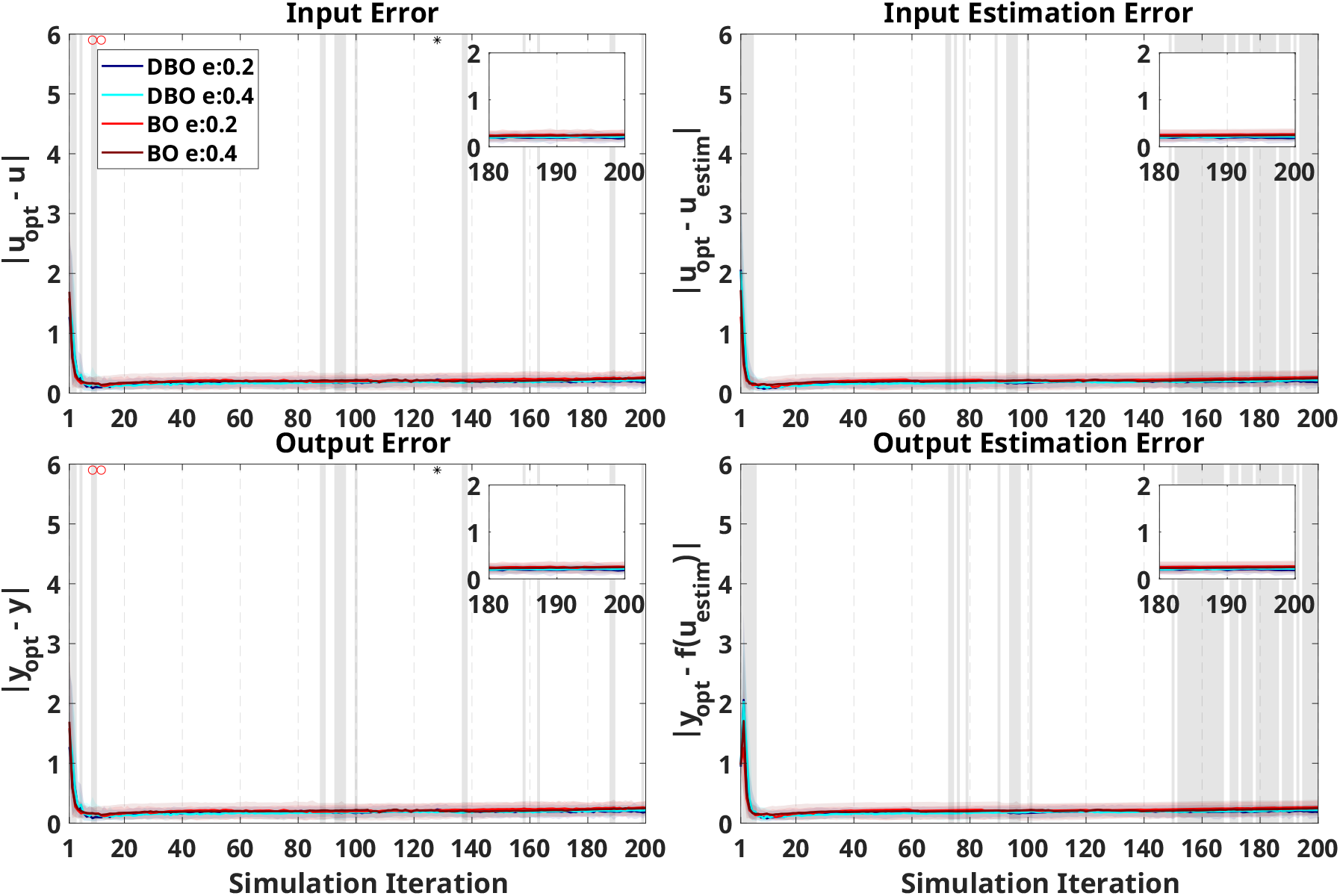
Virtual HIL simulation results using UDL model with slow negative learning (Model 1) as the virtual human response. The *x*-axis indicates the number of iterations during optimization (i.e., after three random inputs are initially applied). In all plots, the blue/cyan and red/crimson indicate simulation results using DBO with e-ratio of 0.2/0.4, and BO with e-ratio of 0.2/0.4, respectively. Solid lines indicate median and shaded regions of same color extend from the 20^th^ to the 80^th^ percentile across 50 repetitions at each iteration. Optimizer performance is quantified based on *input error* (top left), *input estimation error* (top right), *output error* (bottom left), and *output estimation error* (bottom right). An zoomed-in insert is provided in each panel to help visualize differences in the last 20 iterations. The gray shaded region indicates the presence of significant differences (*p*_*unc*_ *<* 0.05) between simulation results using different optimizers at each iteration. Asterisks at the top of each figure indicate a significant effect of the e-ratio, and red circles indicate a significant interaction between e-ratio and the optimizer on the outcome at each iteration.

**Figure S3:**
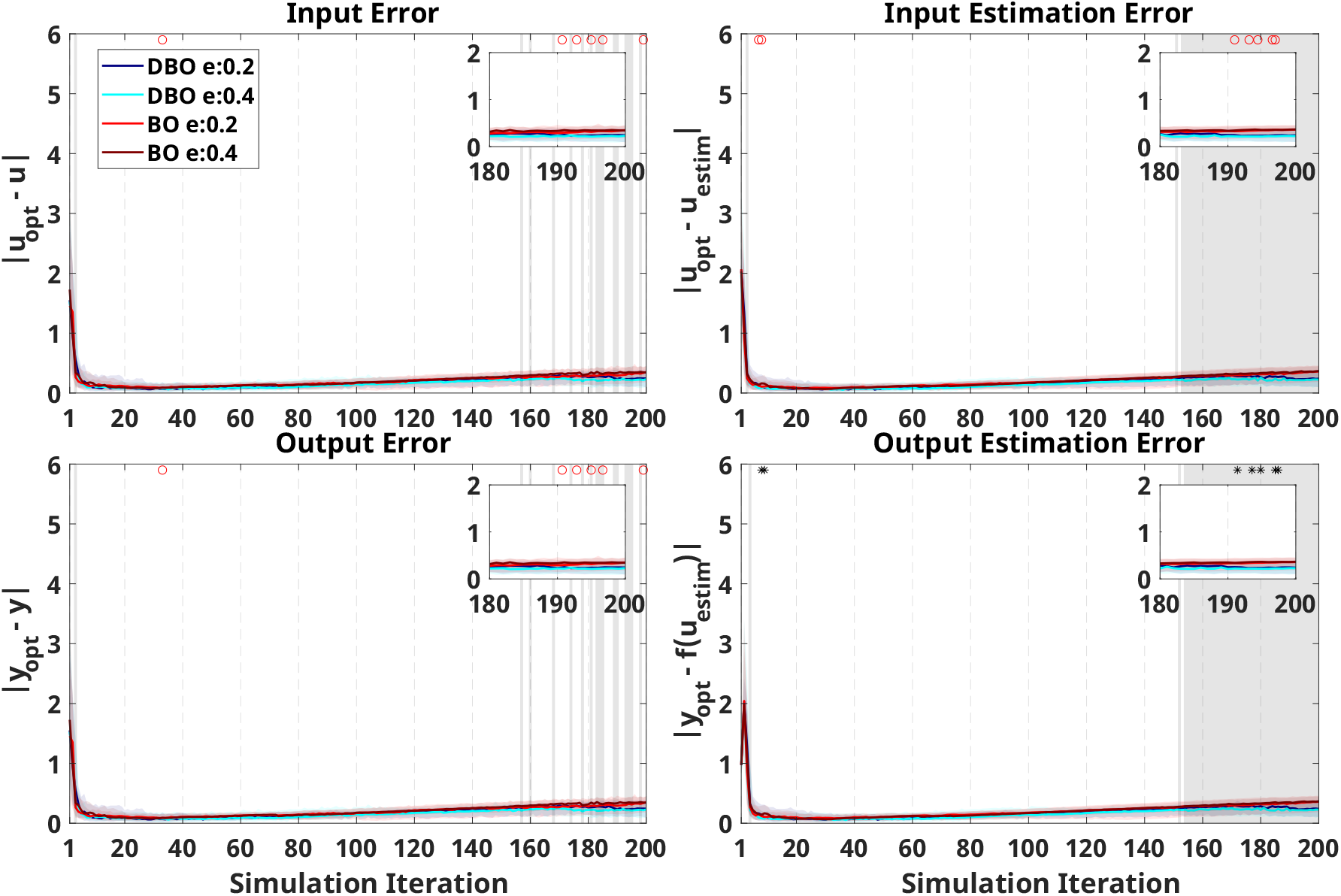
Virtual HIL simulation results using UDL model with slow positive learning (Model 2) as the virtual human response.

**Figure S4:**
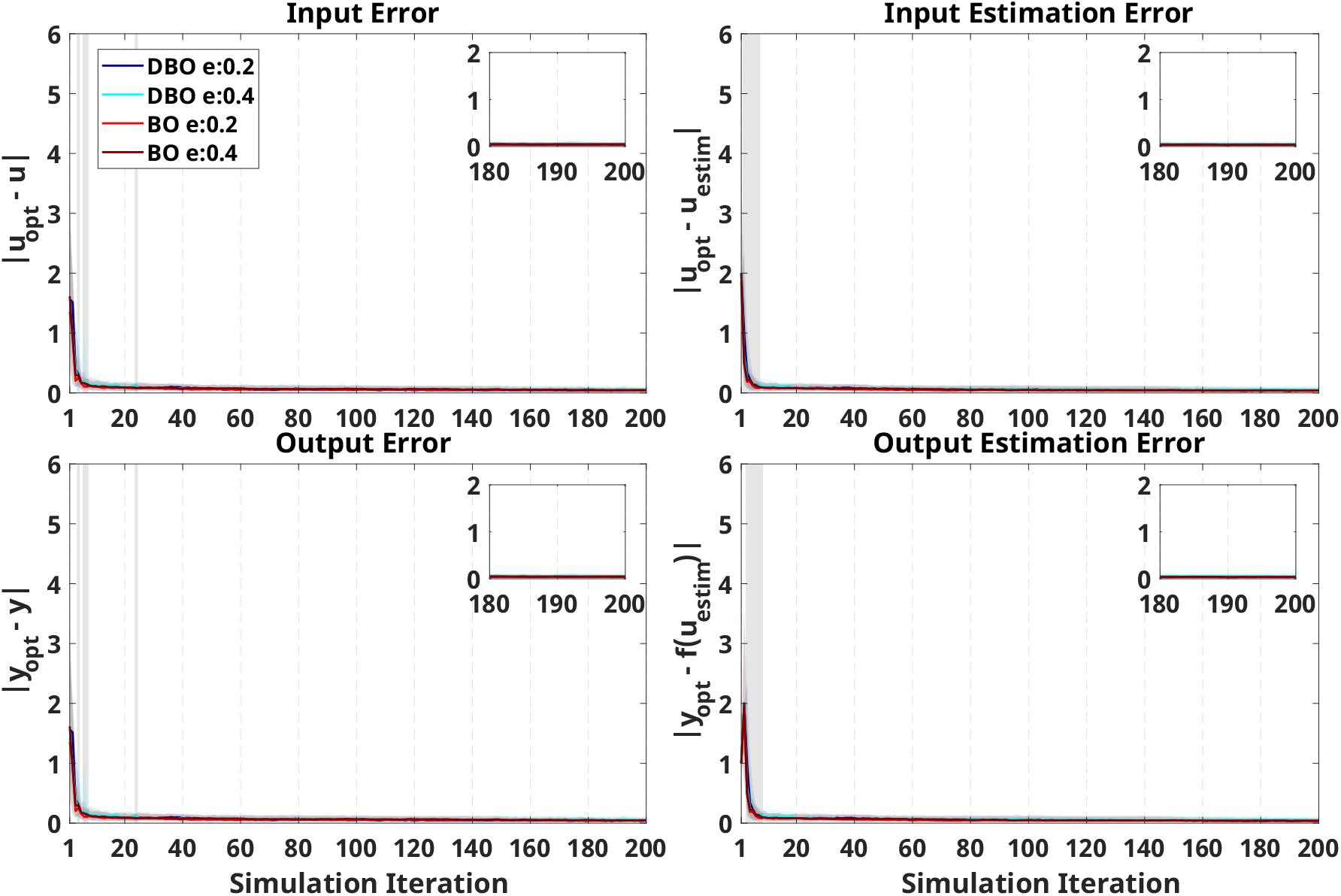
Virtual HIL simulation results using UDL model with constant and no learning (Model 5) as the virtual human response.

**Figure S5:**
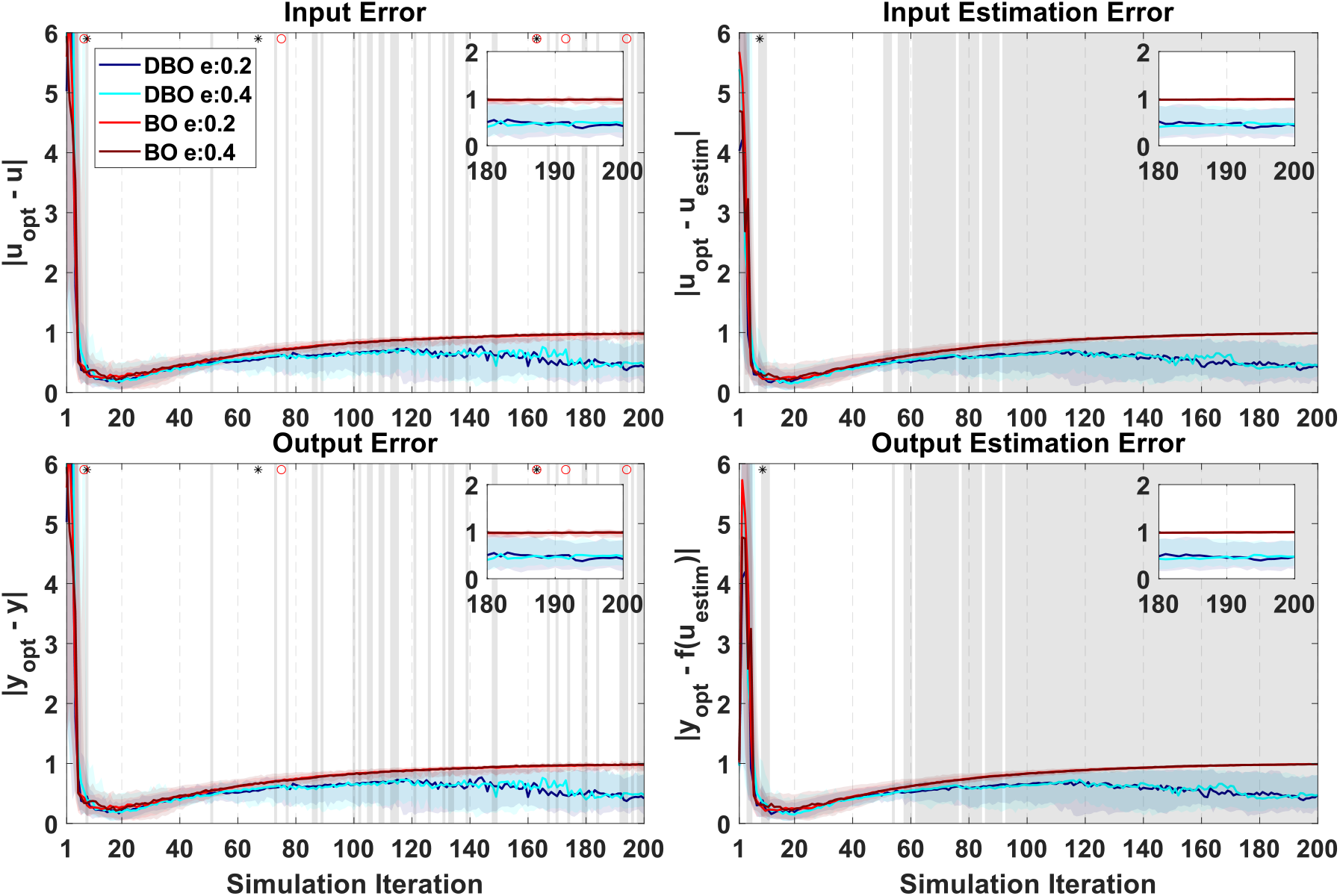
Virtual HIL simulation results using UDL model with only adaptation (Model 6) as the virtual human response.

